# Transposable elements drive species-specific and tissue-specific transcriptomes in human development

**DOI:** 10.1101/2025.05.19.654775

**Authors:** Yun Zhang, Jianqi She, Xueyan Hu, Yueqi Jin, Jiaming Zhao, Songyan Hou, Changyu Tao, Minghao Du, Ence Yang

## Abstract

**Background:** Transposable elements (TEs) are an abundant and crucial regulatory resource in the human genome. Serving as alternative promoters, TEs can be reactivated and produce TE-initiated transcripts, which play importance roles in early development and differentiated tissues. However, the prevalence and function of TE-initiated transcription in human development are poorly characterized.

**Results:** We identified 12,918 TE-initiated transcripts across 40 human body sites and embryonic stem cells. Among TE-initiated transcripts, 80% were activated in a tissue-specific manner. TEs with tissue-specific transcription factor binding motifs were enriched in particular tissues. Additionally, approximately half of TE-derived TSSs were primates-specific. Notably, 375 primates-specific TE-derived TSSs were found to create novel tissue-specific gene expression patterns during evolution.

**Conclusions:** Our results characterize the global profile of TE-initiated transcription and enhance our understanding of TE contribution to the primate-specific gene regulatory networks in human development.

## Background

Transposable elements (TEs) are mobile genetic elements that are widespread in eukaryotic genomes [1,2]. Approximately 45% of the human genome are derived from various classes of TEs, including LINE (long interspersed nuclear element), SINE (short interspersed nuclear element), LTR (long terminal repeat), and DNA transposons [3,4]. With the ability to transpose, TEs have been recognized as a driving force in the evolution of regulatory elements, particularly in bringing about significant changes over short evolutionary time scales [5–10].

As potential threats to human genome, most TEs have been kept inactive by mutational events and epigenetic repression [11,12]. However, growing evidence suggests that certain TEs remain transcriptionally active and play significant roles in embryo development, immune response and tissue homeostasis [13–18]. In our previous study, we reported tissue-specific and biology-associated expression patterns of 13,889 human endogenous retrovirus (HERV, one of the TE classes) RNAs across different human tissues [19]. With abundant regulatory elements, TEs can also function as epigenetically regulated promoters to promote their own transcription or regulate the expression of neighboring genes, producing TE-initiated transcripts with TE-derived transcription start sites (TSSs) [20–22]. Studies have revealed the important roles of TE-initiated transcription in early development and differentiated tissues [14,23,24]. For example, a L1-derived long noncoding RNA (lncRNA), LINC01876, is a human-specific transcript expressed exclusively during brain development, contributing to the differentiation of neural progenitors [25]. In addition, rodent-specific TEs were found to create novel tissue-specific gene expression patterns in mouse by acting as alternative TSSs [6]. Altogether, accumulative evidences suggest a critical need to better understand the function of TEs and their contributions to the evolution of gene regulatory networks in human development.

To address the challenge of ambiguous mapping of short sequencing reads originating from TEs, we incorporated long-read RNA-seq, short-read RNA-seq, CAGE, and RAMPAGE datasets to screen for TE-initiated transcripts. Through this approach, we identified 12,918 TE-initiated transcripts expressed across 40 human body sites and embryonic stem cells. Approximately 80% of these TE-initiated transcripts were tissue-specific expressed, and the activation of TE-derived promoters was strongly associated with tissue-specific epigenetic regulation and transcription factors binding. Using multiz alignment across 470 mammalian genomes, we mapped the evolutionary trajectories of these TE-initiated transcripts and identified 375 primate-specific TE-initiated transcripts that create novel tissue-specific gene expression patterns. Collectively, our findings advance the understanding of how TEs contribute to the primate-specific and tissue-specific gene regulatory networks in human development.

## Results

### Genome-wide identification of TE-initiated transcription in human development

Considering the inherent limitations of TE detection from short reads, we integrated short-read RNA-seq, CAGE, RAMPAGE data with long-read RNA-seq using strict computational framework to screen for TE-initiated transcripts (Fig. 1a). Benefiting from long-read sequencing, our pipeline allowed to accurately capture the features and locations of TEs as well as their connections to downstream genes. For example, the full length of a MLT1K-initiated transcript was identified by long-read RNA-seq, confirming its connection to the TEC gene (Fig. 1b). Compared with TE-initiated transcripts in the TE-TSS database identified by only short-read RNA-seq or CAGE/RAMPAGE data [20], our pipeline showed improved sensitivity and specificity (Fig. 1c; Table S1). Additionally, we performed 5’ rapid amplification of cDNA ends (5’ RACE) and Sanger sequencing for randomly 5 TE-initiated transcripts in K562 cell lines and verified the existence of TE-derived transcription initiation detected by our pipeline (Fig. 1d).

**Fig. 1.**
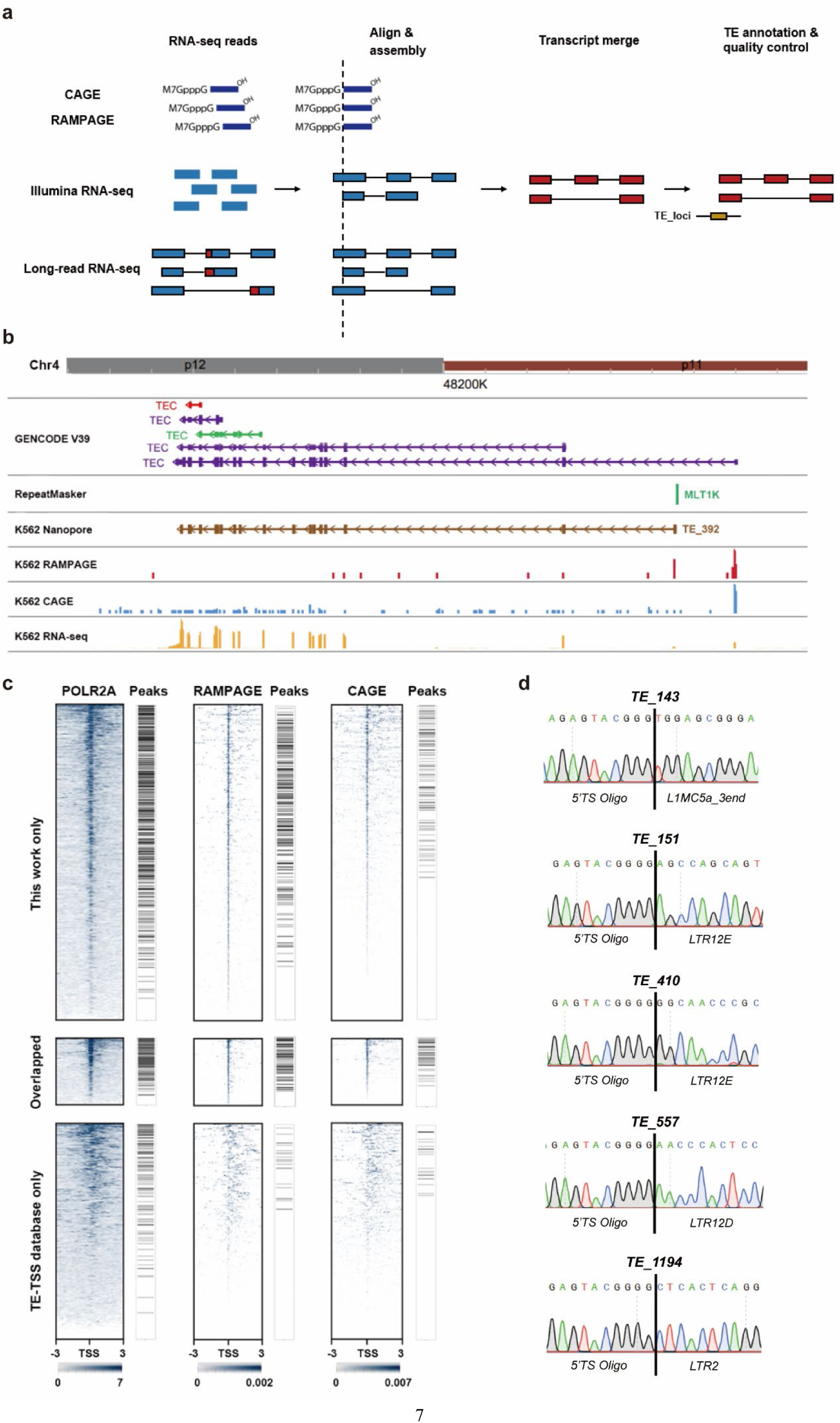
Detection of TE initiated transcripts **a** Pipeline designed to identify TE-initiated transcripts by integrating long-read RNA-seq, short-read RNA-seq, CAGE, and RAMPAGE datasets. **b** A MLT1K-initiated transcript connected with the TEC gene. **c** POLR2A ChIP-seq, RAMPAGE, CAGE signals around the TSSs of TE-initiated transcripts identified in this work, TE-TSS database, or both in K562 cell lines. ChIP-seq peaks are labeled on the right of the corresponding heatmaps. **d** The 5’ RACE sequence results of 5 randomly selected TE-initiated transcripts.

To characterize the landscape of TE-initiated transcriptome and understand the functionality of TEs, we obtained 133 long-read RNA-seq datasets, 9,161 short-read RNA-seq datasets, 69 CAGE datasets and 72 RAMPAGE datasets from GEO, GTEx, FANTOM5 and ENCODE database (Table S2) [26–29]. By analyzing these datasets with our computational pipeline, a total of 12,918 TE-initiated transcripts were identified across 40 human body sites and embryonic stem cells (H1 cell line; Table S3). Annotated with Dfam database [30], we found that these TE transcripts were initiated by 12,749 TE loci from 4 TE classes and 34 TE superfamilies, including LTR (5,331 TE loci from 6 superfamilies), LINE (3,632 TE loci from 6 superfamilies), SINE (2,761 TE loci from 5 superfamilies), and DNA transposons (1,025 TE loci from 17 superfamilies). Enrichment analysis revealed the enrichment of the ERV1, L2 and MaLR superfamilies while the L1 superfamily was conspicuously absent due to widespread 5′ truncations upon insertion [31], which led to the removal of their promoter sequences (Fig. S1). Moreover, these TEs initiated the transcription of 7,458 neighboring genes, including 3,996 protein coding genes and 3,086 lncRNA genes. Interestingly, 2,566 TE-initiated transcripts were predicted to produce novel protein isoforms of protein coding genes, while 542 transcripts connected with lncRNA genes were predicted to have coding potential. Our results highlight that TEs contribute to the generation of abundant transcript and enhance the complexity of transcriptional and translational networks.

### TE-initiated transcription shows high tissue specificity

Previous studies have discovered that TEs are generally expressed with high tissue specificity [6,19,20,32]. In the current work, the majority of TE-initiated transcripts (10,997/12,918) were expressed in a tissue-specific manner, with significant higher Tau index than TE transcripts without TE-TSS (Fig. 2a). The number of TE-initiated transcripts varied across human tissues, ranging from 39 in the minor salivary gland to 6233 in the testis (Fig. 2b). Specially, 92.1% TE-initiated transcripts in testis were exclusively expressed while 74.1% in embryonic cell, suggesting the regulation function of TEs in human germline and development. Additionally, Uniform Manifold Approximation and Projection (UMAP) analysis demonstrated that the expression profiles of TE-initiated transcripts contributed to distinguishing diverse tissues (Fig. 2c).

**Fig. 2.**
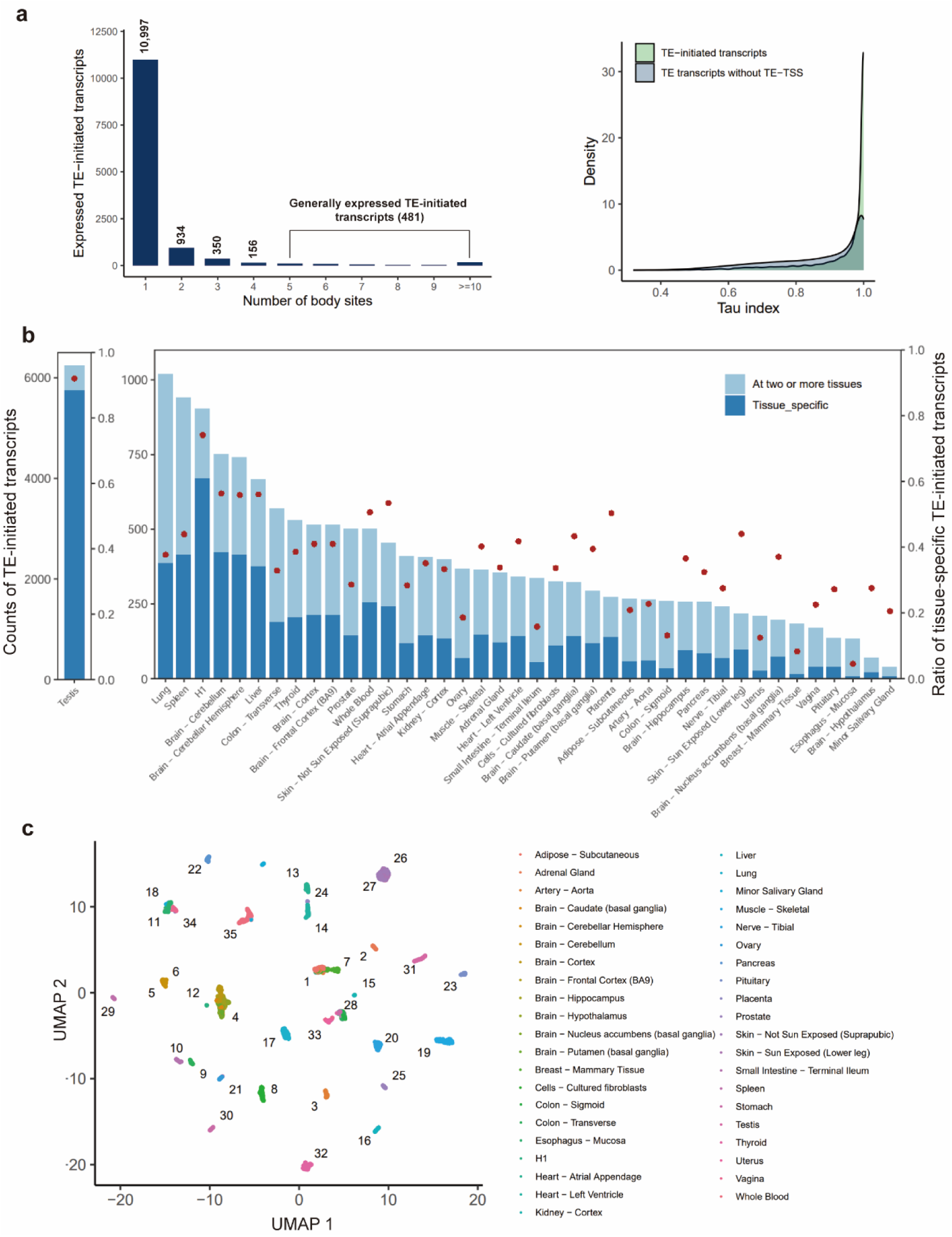
Expressed TE-initiated transcripts show high tissue specificity **a** Histogram showing counts of TE-initiated transcripts expressed in different numbers of body sites (left). Tau index of TE-initiated transcripts and TE transcripts without TE-TSS (right). **b** Distribution of TE-initiated transcripts across 40 body sites and H1 cell line. Blue stacked bars indicate the number of TE-initiated transcripts (left y-axis), and red dots indicate the ratio of tissue-specific TE-initiated transcripts at each body site (right y-axis). **c** Sample similarity based on TE-initiated transcript profiles by UMAP analysis. Tissues are as follows: 1: adipose-subcutaneous; 2: adrenal gland; 3: aorta; 4: brain; 5-6: brain cerebellum; 7: breast; 8: fibroblasts; 9: colon-sigmoid; 10: colon-transverse; 11: esophagus; 12: H1 cell line; 13: heart-atrial appendage; 14: heart-left ventricle; 15: kidney; 16: liver; 17: lung; 18: salivary gland; 19: skeletal muscle; 20: nerve; 21:ovary; 22: pancreas; 23: pituitary; 24: placenta; 25: prostate; 26-27: skin; 28: small intestine; 29: spleen; 30: stomach; 31: testis; 32: thyroid; 33: uterus; 34: vagina; 35: whole blood.

Given that TEs are considered as abundant gene regulators in the human genome [31], we then investigated whether TE-initiated transcription influenced the tissue specificity of gene expression. Compared with genes without TE-derived TSSs, TE-initiated genes that used TE-derived TSSs more often showed significant higher tissue specificity (Fig. 3a). For example, MER61D was found to initiate the liver-specific transcription of CYP2C18 (Fig. 3b), a cytochrome P450 protein involved in liver drug metabolism [33]. For embryonic cells, TE-initiated genes were significantly enriched in stemness gene signatures [34], including embryonic stem expressed genes and Nanog targets, suggesting that TEs may play a crucial role in human development by interacting with Nanog (Fig. 3c). Through gene function enrichment analysis for each tissue, our results showed that TE-initiated genes were associated with tissue-relevant function and pathways (Fig. S2; Table S4). For example, TE-initiated genes identified in the kidney cortex were linked to drug ADME (Fig. 3d).

**Fig. 3.**
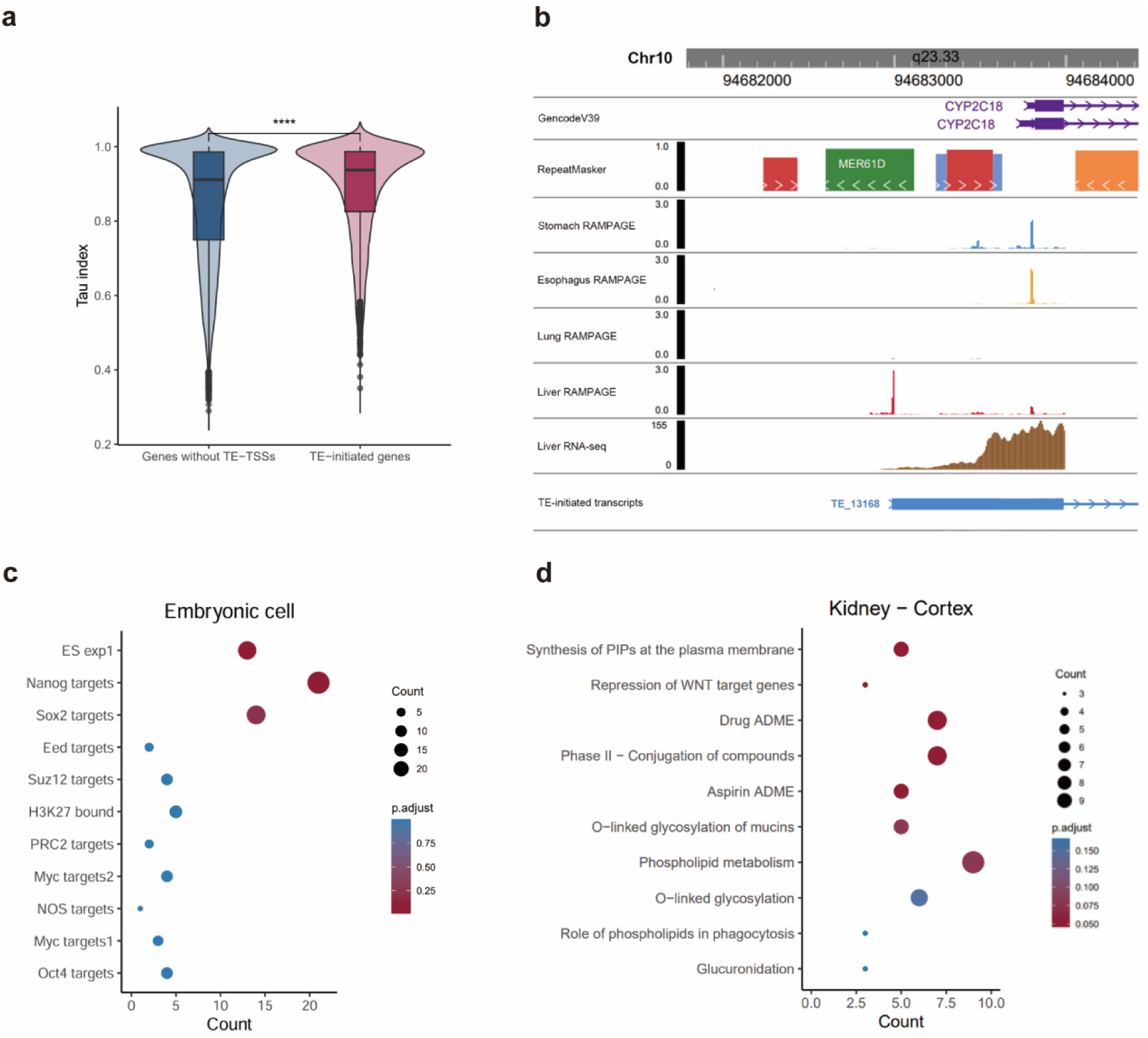
TEs can initiate the tissue-specific transcription of genes **a** Tau index of TE-initiated genes (used TE-derived TSSs more often) and genes without TE-TSS **b** Browser view of RAMPAGE signals and RNA-seq coverage around the TSS of TE_13168. **c** The enrichment analysis of stemness gene signatures for TE-initiated genes in embryonic cells. **d** The reactome enrichment analysis of TE-initiated genes in kidney cortex. The size of point represents the count of TE-initiated genes, and the gradual color change from red to blue represents the adjust *P* value change from low to high.

### Tissue-specific TE-initiated transcription were associated with tissue-specific transcription factors

To further understand the tissue specificity of TE-initiated transcription, we firstly investigated the sequence determinants of these TEs. By TE family enrichment analysis, we identified 62 TE families significantly overrepresenting in specific tissues (Fig. 4a and S3; Table S5). The LTR12C, LTR12D and LTR12E families, which were reported to exhibit strong regulatory ability in human adult spermatogonial stem cells (hSSCs) [35], were particularly enriched in the testis (Fig. 4b). The LTR7, L1HS_5end, and HERVH families, biomarkers for preimplantation embryo development [23], were significantly enriched in the embryonic cell (Fig. 4b). Next, we asked whether these enriched TE families share certain tissue-specific transcription factor binding motifs and have been co-opted as tissue-specific promoters. With transcription factor enrichment analysis, we identified 44/62 TE families harboring tissue-specific transcription factor binding sites (TFBS) (Fig. 4c and S4; Table S6). For example, the LTR7 family was significantly enriched in the binding sites of NANOG as previously reported (Fig. 4d) [36], which is involved in embryonic stem cell proliferation, renewal, and pluripotency [37]. The skin-overrepresented HERVE family was enriched in the binding sites of TFAP2C (Fig. 4d), which is associated with the keratinocyte maturation [38].

**Fig. 4.**
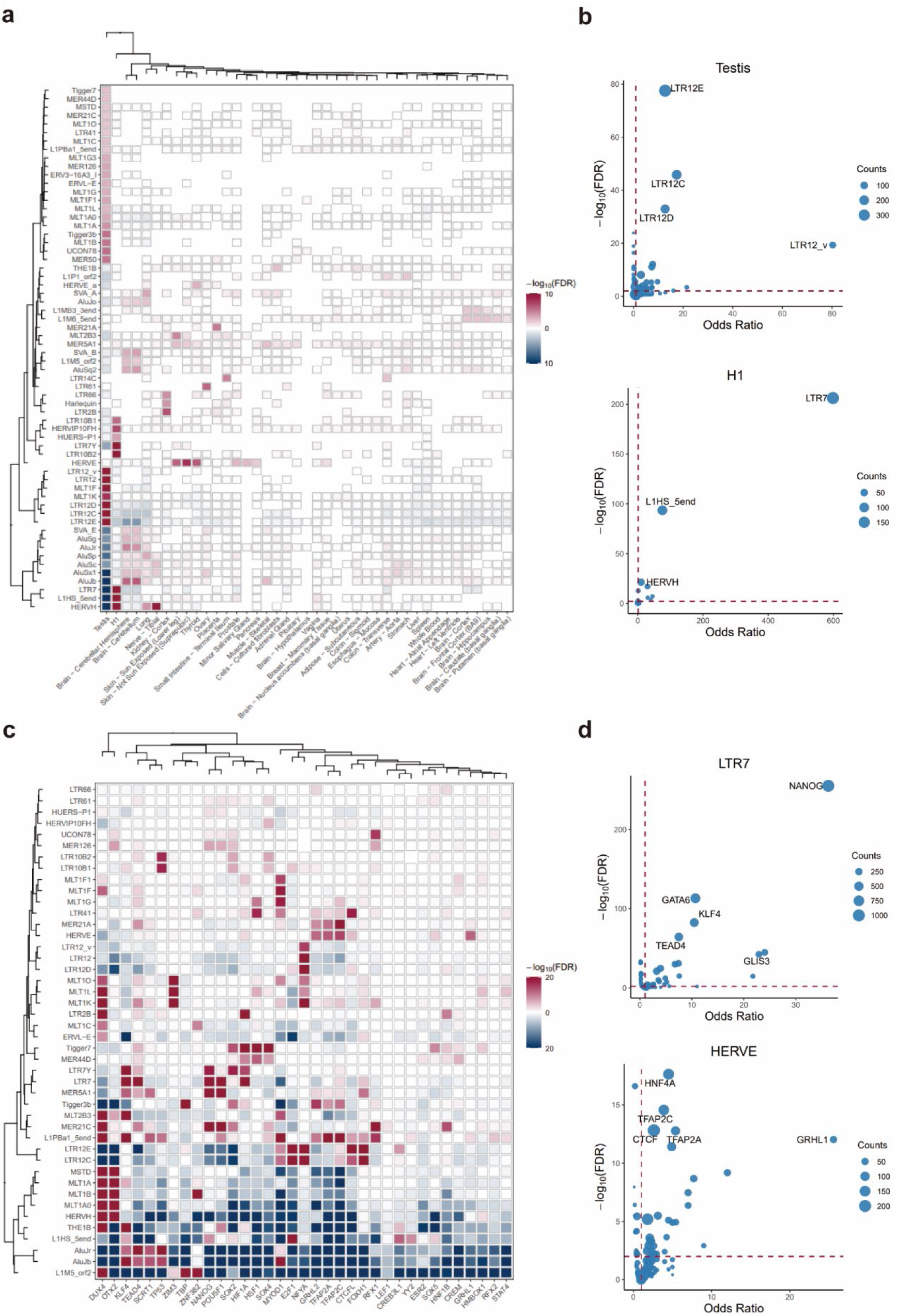
The transcription factor binding contributes to the tissue-specific expression of TE-initiated transcripts. **a** The enrichment analysis of TE families for TE-initiated transcripts across various tissues is displayed using color intensities corresponding to their-log_10_(FDR) values. Red indicates overrepresentation, while blue indicates depletion. **b** The enrichment analysis of TE families for TE-initiated transcripts in testis and embryonic cells. The size of point represents the count of TE-initiated transcripts. **c** The enrichment analysis of TFBS for 44 TE families. **d** The enrichment analysis of TFBS in LTR7 and HERVE family. The size of point represents the count of TE loci.

Since TE expression is generally suppressed by DNA methylation and/or repressive histone modifications [11,21], we also investigated epigenetic events of TE-initiated transcription from ENTEx datasets [26]. As expected, the DNA methylation level of TE-derived TSSs were significantly lower than that of silent TEs in all tissues examined (Fig. 5a). Moreover, the DNA methylation level of tissue-specific TE-derived TSSs in the specific tissue were significantly lower than that in other tissues, suggesting that TE promoters are activated by DNA demethylation in a tissue-specific manner (Fig. 5b and S5). Additionally, active histone modifications, such as H3K27 acetylation, H3K4 monomethylation and H3K4 trimethylation, were significantly enriched within ±1 kb of tissue-specific TE-derived TSSs (Fig. 5c). Consequently, the epigenetic status supported the conclusion that the regulatory potential of these TE-initiated transcription is unlocked and reactivated in specific tissues.

**Fig. 5.**
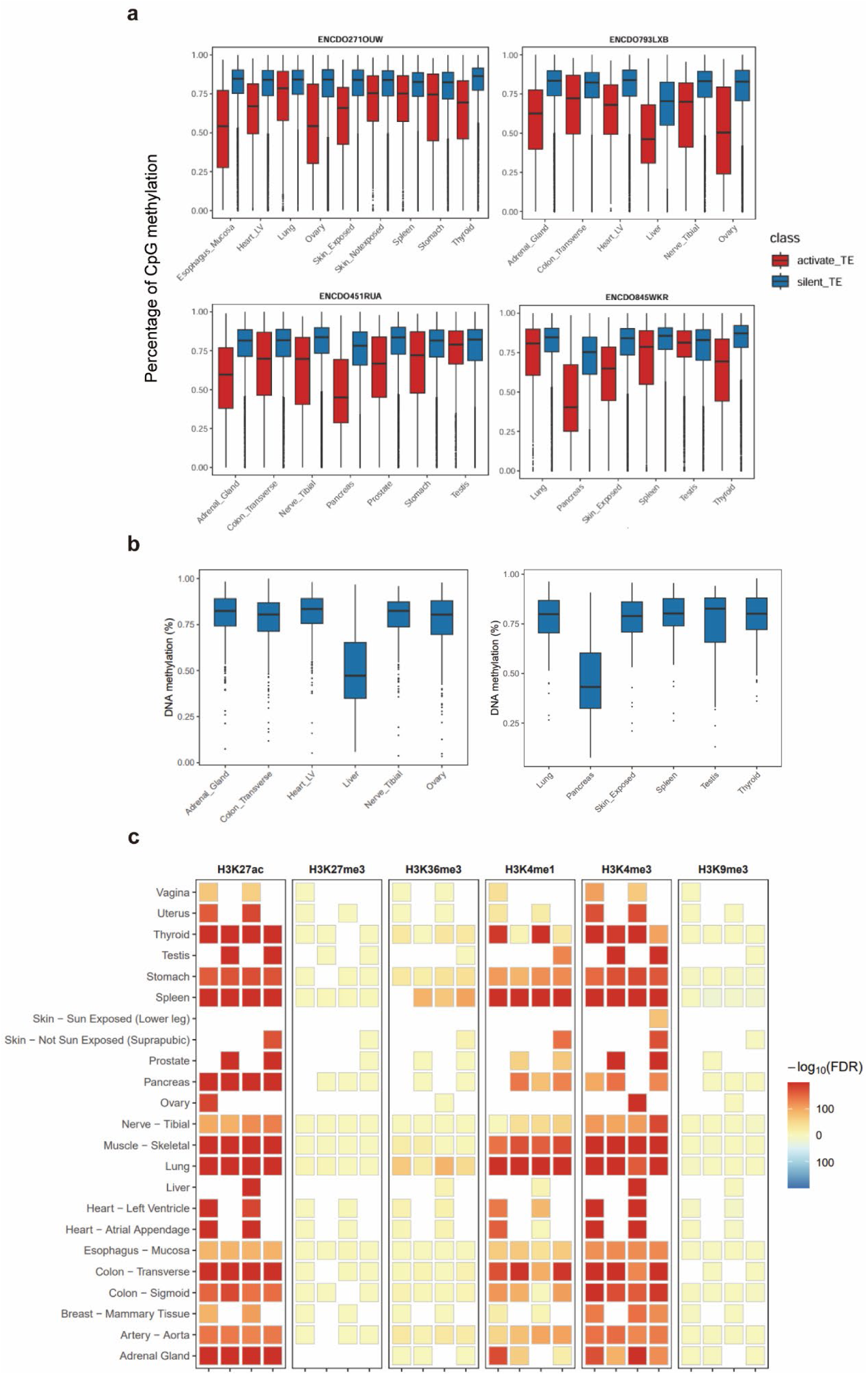
Tissue-specific TE-initiated transcripts expression corresponds to tissue-specific epigenetic regulation **a** The boxplot shows the distribution of the DNA methylation levels of expressed TE-initiated transcripts and silent TE-initiated transcripts at each body site. The difference between the expressed TE-initiated transcripts and silent TE-initiated transcripts at each body site was significant (*P* < 0.01, Wilcoxon rank-sum test). **b** The boxplot shows the distribution of the DNA methylation levels of tissue-specific TE-initiated transcripts in the specific tissue and other tissues (left: liver; right: pancreas). The difference between the specific tissue and other tissues was significant (*P* < 0.01, Wilcoxon rank-sum test). **c** The enrichment analyses of histone modifications for expressed TE-initiated transcripts within each body site in each individual (from left to right: ENCDO271OUW, ENCDO451RUA, ENCDO793LXB, ENCDO845WKR).

### TE-derived TSSs with distinct patterns of evolutionary conservation

Next, we examined the evolutionary conservation of TE-derived TSSs identified in our study. Using multiz alignment across 470 mammalian genomes [5,39], we found 88.5% of all human TEs and 95.4% of TE-derived TSSs had orthologous counterparts, while 98.1% of traditional TSS regions without TEs had orthologous sequences. We categorized 12,918 TE-derived TSSs into 3 groups with distinct evolutionary patterns: (1) Group 1 (G1) for highly conserved TE-derived TSSs, aligned with ≥ 50% to over half of the mammalian genomes; (2) Group 2 (G2) for primate-specific TE-derived TSSs, with ≥ 50% alignment limited to primate genomes; (3) Group 3 (G3) for other evolving TE-derived TSSs (Fig. 6a and b, Table S7). Compared with traditional TSS regions from protein coding genes and lncRNA genes, TE-derived TSSs have the lowest proportion of G1 (41.9%) but highest proportion of G2 (46.0%) (Fig. 6b). Among these TE-derived TSSs, LTRs have the lowest proportion in G1 (26.4%, *P* < 2.2 × 10^-16^ compared with randomly sampled genomic regions) and the highest proportion in G2 (61.7%, *P* < 2.2 × 10^-16^). By analyzing the 470-ways PhastCons conservation [39], we also found that TE-derived TSSs showed much lower PhastCons scores than traditional TSS regions from protein coding genes and lncRNA genes (Fig. 6c). These results suggested that TEs are significant driving forces in the evolution of regulatory elements, creating novel TSSs that initiate transcription in primate genomes.

**Fig. 6.**
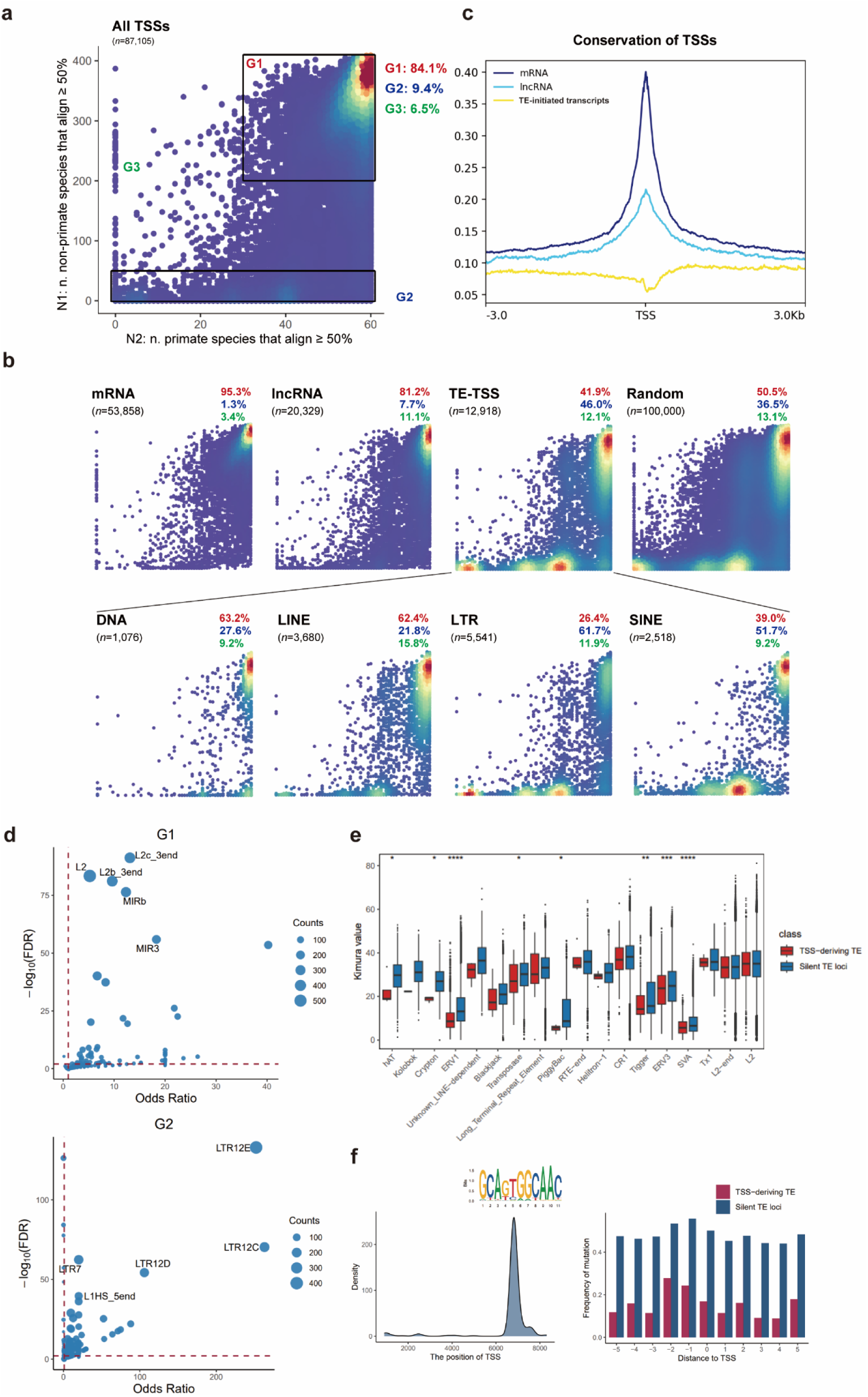
Subsets of TSS regions show distinct patterns of evolutionary conservation. **a** Distribution of human TSS regions according to the genomes in which they align. N1 represents the number of non-primate species in which ≥ 50% of the nucleotides align. N2 represents the number of primate species in which ≥ 50% of the nucleotides align. Three groups, corresponding to dense regions in the heatmap, are highlighted. **b** As in (a) but illustrating distributions of TSS regions of TE-initiated transcripts, mRNAs, lncRNAs, random regions, and different classes of TE-initiated transcripts; fractions of TSS regions in each of the three groups mentioned in a are indicated. **c** Averaged PhastCons score at 6-kb regions centered by TSSs of TE-initiated transcripts, mRNAs, and lncRNAs. TSSs of TE-initiated transcripts showed the lowest phastCons score in the center. **d** The TE family enrichment analysis of different groups of TE-initiated transcripts. **e** The boxplot shows the distribution of the Kimura values of TSS-deriving TEs and silent TE loci for 18 TE families. **f** The density of TSSs within the consensus sequence of LTR12E (left). The mutation frequency of TSS motif in TSS-deriving TEs and silent TE loci (right).

To investigate the features of TE-derived TSSs with distinct patterns of evolutionary conservation, we firstly performed TE family enrichment analysis on each group of the TE-TSSs (Table S8). G1 TE-derived TSSs were significantly enriched in the older TE families from L2, Charlie and CR1, while G2 TE-derived TSSs were enriched in the younger TE families from ERV1. The top 5 TE families represented in the G2 group, including LTR12E, LTR12C, LTR7, LTR12D, and L1HS_5end were all previously recognized as primate-specific TE families (Fig. 6d). We further compared the TE evolutionary age between TSS-deriving TEs and silent TE loci. Notably, younger TEs were particularly inclined toward active transcription in 8 out of 34 TE superfamilies, especially the ERV1 superfamily (Fig. 6e). Moreover, compared with silent LTR12E loci, the mutation frequency of the TSS motif in TSS-deriving LTR12Es were significantly lower (Fig. 6f), which suggested that recently inserted TEs, which experience lower negative selective pressure or undergo positive selection, retain primary sequences for active transcription.

Besides TE-derived TSSs, their downstream genes also exhibited distinct evolutionary pattern. Compared to G1 conserved TE-TSSs, G2 primate-specific TE-TSSs initiated the transcription of a higher proportion of primate-specific protein-coding genes (54/1577, *P* = 2.4 × 10^-10^) and lncRNA genes (33/458, *P* = 1.2 × 10^-4^). TE-initiated genes of G1 were most enriched in the RHO GTPase cycle (FDR = 7.8 × 10^-4^, Fig. S6 and Table S9), an essential biological process. Conversely, TE-initiated genes of G2 were enriched in biological processes involving interaction with the environment, such as drug ADME (FDR = 0.053, Fig. S6). These results suggested that primate-specific TEs contributed to the origination of lineage-specific genes during evolution and adaption to environment.

### Primate-specific TEs altered the tissue specificity of gene expression

Previous study has demonstrated the contribution of rodent-specific TEs to creating novel tissue-specific gene expression patterns by acting as alternative TSSs [6]. To ask whether primate-specific TEs have similar impact, we compared the tissue expression patterns between human and mouse for 1,144 homologous genes associated with 1,308 primate-specific TE-TSSs. Utilizing gene expression data of 17 tissues obtained from the GTEx and ENCODE datasets [26,27], we found that 37.5% homologous genes exhibited distinct expression patterns at the tissue level. Notably, 375 primate-specific TE-derived TSSs were identified to create novel tissue-specific gene expression in humans (*Z* score difference > 1, Fig. 7a, Table S10) [40]. For example, IL27 encodes one of the subunits of a heterodimeric cytokine complex, which could drive rapid expansion of naive CD4(+) T cells and promote adipocyte thermogenesis and energy expenditure [41]. In the human genome, we identified a MER67A-derived TSS at the 19.5 kb upstream from the canonical TSS of IL27 (Fig. 7b). The integration of MER67A created a stronger TSS to initiate the transcription of IL27 and we found that IL27 exhibited the highest expression level in the human liver. However, in the all corresponding 16 mouse tissues (including liver), IL27 showed significant low expression level, possibly due to the ability of MER67A to respond to liver-specific transcription factors, including FOXA1, FOXA2, HNF4A, and HNF4G (Fig. 7b and c). Additionally, two novel transcription factor motifs, HNF4A and SP1, identified using FIMO [42], were generated by mutations specific to the MER67A promoter of IL27 (Fig. 7d). These findings suggested that domesticated TEs could alter the tissue specificity of gene expression as primate-specific alternative promoters.

**Fig. 7.**
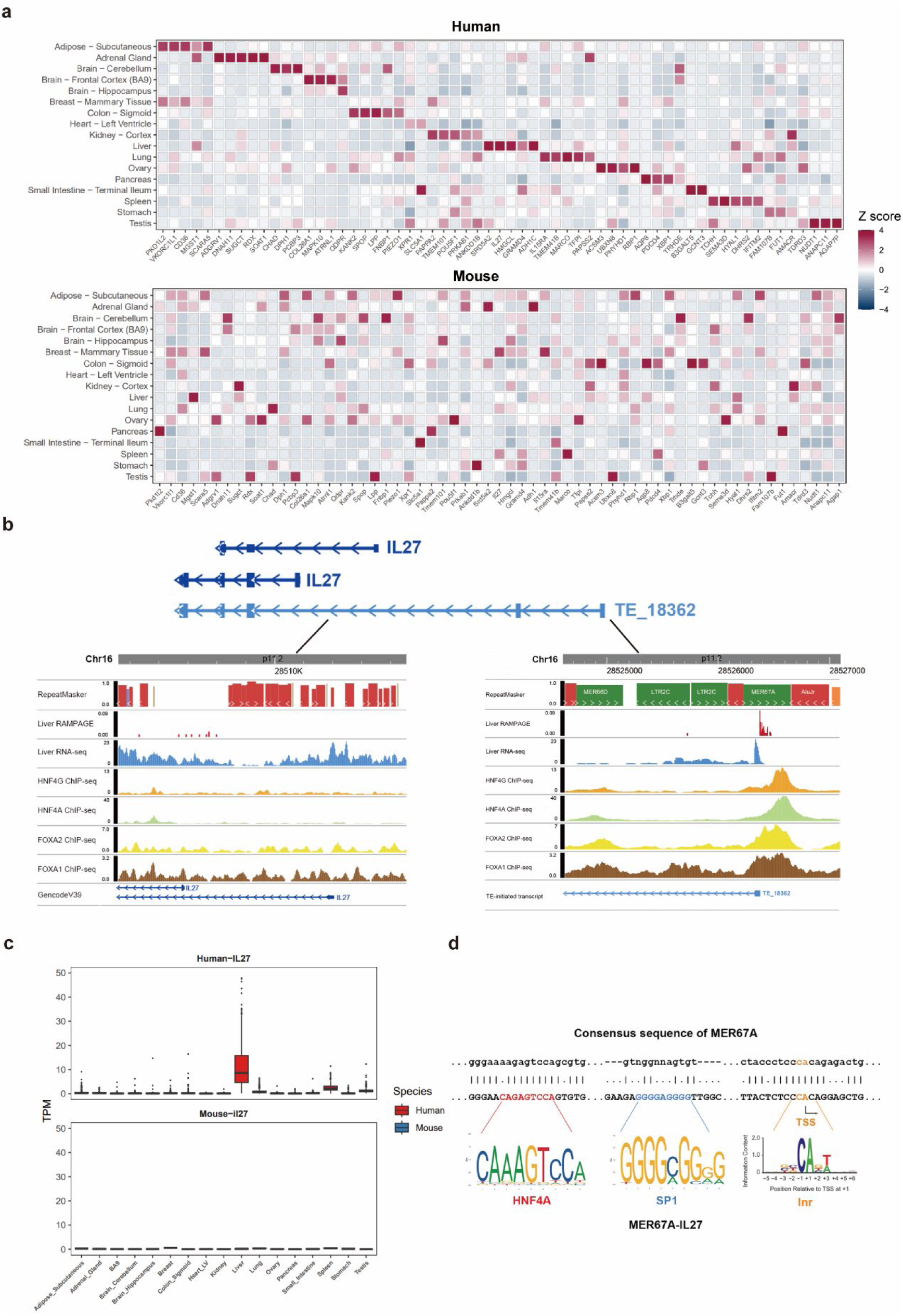
Primates-specific TE drive the expression of tissue-specific genes. **a** Expression pattern of homologous protein-coding genes associated with primate-specific TE-initiated transcripts in 17 tissues of mouse and human. **b** Browser view of RAMPAGE signals, RNA-seq coverage, and liver-specific transcription factors ChIP-seq signals around the TSS of TE_18362 and canonical TSSs of IL27. **c** Expression pattern of IL27 in 17 tissues of mouse and human. **d** The comparison between the consensus sequence of MER67A and the MER67A promoter of IL27.

## Discussion

As “genomic parasites”, TEs often carry sequences that recruit host transcription machinery, facilitating their own transcription and transposition [43]. Although most TEs have lost their ability to transpose, some TEs can be domesticated as regulatory elements, such as alternative promoters, which promote TE-initiated transcription and play vital roles in human development as well as the evolution of gene regulation [6,14,20,24,44,45]. In this study, we identified 12,918 TE-initiated transcripts across human body sites and embryonic stem cells, exhibiting species-specific and tissue-specific expression pattern in human development. Moreover, 5,262 TE-initiated transcripts have protein-coding potential and even produce novel protein isoforms. Overall, these abundant TE-initiated transcripts contribute to gene expression regulation and the generation of diverse protein isoforms, suggesting that some TEs escape epigenetic silencing and have been co-opted as alternative promoters, benefiting the host.

TE-initiated transcripts have recently been identified using transcript assembly from short-read RNA-seq data [20,22]. However, for short-read sequencing, the RNA molecules are fragmented, often leading to the loss of TSS information [46]. Techniques like CAGE and RAMPAGE enable accurate TSS identification at base-pair resolution but fail to capture the full length of TE-initiated transcripts [28,47]. Long-read sequencing technologies, such as Pacific Biosciences (PacBio) and Oxford Nanopore Technologies (ONT), offer the advantage of generating longer reads (>10 kb), enabling the identification of full-length isoforms [48,49]. However, these technologies are not able to always guarantee the 5’ completeness of the sequenced product, with a significant proportion of cDNA 5’ ends falling short of actual TSSs [50]. To overcome these limitations, our study developed a pipeline by integrating long-read RNA-seq, short-read RNA-seq, CAGE, and RAMPAGE datasets to detect TE-initiated transcripts. Benefiting from multi-technologies correction, we successfully captured both accurate TSSs and full-length transcripts, which contributed to deciphering the expression pattern and function of TEs. Due to the imperfect matching between TSSs and full-length transcripts, the identification of TE-initiated transcripts was also not flawless. Combining the Cap-trapping strategy with long-read sequencing technologies, such as CapTrap-seq [51], may offer an improved approach.

Previous studies have shown that TEs can spread binding sites for multiple transcription factors, including CTCF, TP53, ESR1, POU5F1, SOX2, STAT1, and NANOG [7,52,53]. In this study, we observed that TEs with tissue-specific transcription factor binding motifs were enriched in particular tissues. For example, the HERVE family was enriched with TFAP2C binding sites, and HERVE-initiated transcripts were specifically expressed in sun-exposed skin. TFAP2C is known to drive surface ectoderm differentiation, and progenitor cells initiated by TFAP2C can mature into functional keratinocytes [38]. In another case, the HERVH family was enriched with GATA6 binding sites (Fig. S4), and TE-initiated transcripts from HERVH were specifically expressed in the lung and tibial nerve (Fig. S3). GATA6 is known to be required for lung maturation and its inhibition will block terminal differentiation. The results suggest that the activation of TE promoters may play a crucial role in the gene regulatory networks involved in early tissue development. However, whether the TE promoters are activated during the keratinocyte maturation and lung epithelial cell differentiation requires further investigation using additional experimental approaches, such as single-cell RNA sequencing.

We also found that half of the TE-initiated transcripts were primate-specific, with their TSSs exhibiting significantly lower PhastCons scores compared to mRNAs and lncRNAs. This finding underscores the critical role of TEs in the evolution of primate-specific regulatory elements. As alternative promoters, TEs can drive novel tissue-specific gene expression patterns, providing a unique mechanism to accelerate regulatory evolution in species they invade. For example, the integration of the MER67A created a novel TSS for IL27, with subsequent mutations forming a liver-specific HNF4A binding site that initiated liver-specific expression of a MER67A-initiated IL27 transcript. Similarly, the integration of the LTR2B created a novel TSS for UGT2B7 in the human kidney, resulting in a chimeric truncated protein isoform, potentially due to sequence features of LTR2B responsive to the kidney-specific transcription factor HNF1A (Fig. S4). Our results illustrate how TE insertions facilitate the establishment of tissue-specific promoters, thereby expediting regulatory evolution.

## Conclusions

In summary, our work identified over ten thousand TE-initiated transcripts across various human body sites. Most of these TE-initiated transcripts show high tissue specificity and are associated with the spread of tissue-specific TFBSs by TEs. TEs contributed to the emergence of many novel regulatory elements in the primate lineage. Our study also found that half of these TE-initiated transcripts were primate-specific and some of them could create novel tissue-specific gene expression patterns in human. Overall, our study provides a comprehensive investigation into the role of TEs in shaping tissue-specific gene regulatory networks of primates.

## Methods

### Data download

We collected 9,175 short-read RNA-seq samples from several public databases, including the GTEx (by dbGaP, study accession: phs000424.v8.p2), ENCODE, and HPA project [54]. Long-read RNA-seq datasets were download from the GTEx, ENCODE, and the Gene Expression Omnibus database (GSE192955) [29]. CAGE datasets were download from the Fantom5 project (https://fantom.gsc.riken.jp/5/datafiles/latest/). RAMPAGE datasets, ChIP-seq datasets, and epigenomic processed data were download from the ENCODE project. Essential meta-information, such as biosample characteristics and sequencing types, was systematically gathered (Table S1). GENCODE Version 44 was download from https://www.gencodegenes.org/human/ and used as the transcript reference. TE annotations and consensus sequences of TE family were downloaded from the Dfam database (https://www.dfam.org/releases/Dfam_3.8/). The 470-way multiz alignment and 470-mammal phastCons scores were downloaded from the UCSC (https://hgdownload.soe.ucsc.edu/goldenPath/hg38/).

### Pipeline for the detection of TE initiated RNAs

First, RNA-seq data were aligned to the reference human genome (GRCh38) using STAR v2.7.10b and minimap2 [55,56]. For short-read RNA-seq samples, StringTie v2.1.7 was used to assemble the reads into full-length transcripts (stringtie –m 200 –c 1-t) [57]. For long-read RNA-seq samples, FLAIR v.1.4 was employed to correct misaligned splice sites according to the GENCODE v44 and short-read splice junctions [58,59]. For CAGE and RAMPAGE, BEDTools v2.31.1 [60] was used to extract the counts of transcription start sites (TSSs). These datasets were then integrated with a custom script. Briefly, the first exons of multi-exon transcripts, including those generated from the StringTie assembly, FLAIR correction, and GENCODE v44, were extracted. Following this, first exons overlapping with other internal exons were excluded, as they may have been generated from RNA degradation. Then, for each first exon, the TSS was corrected within a range of 20 to 2,588 bp (the 99th percentile of all GENCODE v44 transcript first exons) upstream of its 3’ end, using weight scores [59] calculated from integrated datasets as follows:

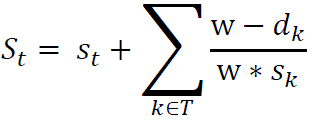

*S* is the weight score for position *t*; *s* represents the counts of reads (including long-read RNA-seq, CAGE, and RAMPAGE) supporting the position as a TSS; *w* is the window size used to calculate the weight score; *T* is the set of all positions within a distance from *t* that is less than the window size; and *d* is the distance between positions *t* and *k*.

For single-exon transcripts, most steps were similar; however, strand orientation was corrected based on overlapping TEs. Corrected TSSs overlapping with TE were retained, and the full-length transcripts of TE-initiated transcripts were merged using reference-guided approach across samples from the same body site. Subsequently, TE-initiated transcripts were consolidated across distinct body sites. Finally, the expression of TE-initiated transcripts was calculated at the transcript level using StringTie. TE-initiated transcripts with a TPM > 0 in at least 50% of samples from the same body site were considered expressed.

### Computational validation of TE-initiated transcripts identified in K562

To evaluate the precision of expressed TE-initiated transcripts identified by our pipeline, we downloaded the ChIP-seq profiles of POLR2A, CAGE and RAMPAGE datasets in K562 cells from the ENCODE project. Then the plots of the POLR2A binding, CAGE signals and RAMPAGE signals surrounding the TSSs were generated using deepTools [61]. The example of TE-initiated transcripts was visualized on WashU Epigenome Browser [62].

### Cell culture and experimental validation of TE-initiated transcripts

The K562 cell line were cultured under 37 °C/5% CO_2_ conditions in complete medium. RNA was isolated from cell pellets containing 5 × 10^6^ cells using the FastPure® Cell/Tissue Total RNA Isolation Kit V2 (Vazyme). The isolated total RNA was applied for cloning the 5′ cDNA end sequences by a HiScript-TS 5′ RACE Kit (Vazyme) according to the protocol provided in the kit with specific primers (Table S11). The amplified fragment was identified by Sanger sequencing.

### Annotation and expression analysis of TE-initiated transcripts

Identified TE-initiated transcripts were annotated with Dfam database [30] by using BEDTools v2.31.1. The relationship between TE-initiated transcript and the closest reference transcript were classified using GffCompare v0.11.2 [63]. The transcripts classified as “ckmnje” were merged into the chimeric TE-initiated transcripts. Coding potential of TE-initiated transcripts was calculated by CPC2 [64]. For coding TE-initiated transcripts, we subsequently compared predicted CDS with annotated CDS using BLAST 2.14.1 [65]. Then the coding TE-initiated transcripts were annotated into five classes [22]: annotated, 5’ chimeric normal, 5’ chimeric truncated, 5’ truncated and novel. The UMAP analysis was conducted using uwot package [66]. The tissue specificity was evaluated with a previously proposed index, which varies between 0 for housekeeping genes and 1 for tissue-specific genes [67]. The reactome pathway enrichment analysis of gene-chimeric TE-initiated transcripts was conducted by the ReactomePA package [68].

### Enrichment analysis of TE family and TFBS

For each TE family, we calculated the number of expressed TE-initiated transcripts (identified in the body site) and silent TE-initiated transcripts (identified in other body sites), both associated with and not associated with the TE family. Then, a two-tailed Fisher’s exact test was applied to assess the enrichment of TE family. For the enrichment analyses of TFBS, we obtained the chromosome coordinates of TFBSs for 367 TFs from the GTRD database [5]. For each TE family, background region sets were generated by randomly selecting regions based on TE annotations using BEDTools v2.31.1. The enrichment of TFBS relative to the random regions was then assessed using a two-tailed Fisher’s exact test. All *P*-values were corrected using the Benjamini-Hochberg (BH) procedure.

### Epigenetic regulation analysis of TE-initiated transcripts

For epigenomic data from the ENTEx Project [26], we calculated the DNA methylation level and extracted histone peaks within ±1 kb of the TSSs of TE-initiated transcripts at all body sites. For each sample, we compared the DNA methylation levels between identified TE-initiated transcripts and silent TE-initiated transcripts (identified in other tissues) using a two-sided Wilcoxon rank-sum test. The enrichment analyses of histone modifications were performed using the same approach as the TE family enrichment analysis. Regions ±1 kb from the TSSs of silent TE-initiated transcripts were used as background sets.

### Estimating the evolutionary conservation of TE-initiated transcripts

To identify groups of TSSs with distinct evolutionary conservation patterns, we computed N1 and N2—the number of non-primate and primate genomes that could align with ≥ 50% of the region’s nucleotides aligned with humans. Three distinct conservation groups emerged: Group 1: highly conserved (N1 ≥ 200 and N2 ≥ 30); Group 2: primate-specific (N1 ≤ 50); and Group 3: other actively evolving. Additionally, the plots of the phastCons scores surrounding the TSSs were generated using deepTools [61]. The Kimura value of TE loci was calculated using RepeatMasker [30]. The flanking sequences of LTR12E-derived TSSs and the DNA sequences of silent LTR12E loci (no associated TE-initiated transcripts identified) were aligned using MAFFT with default parameters [69].

### The comparison of tissue expression patterns for homologous genes

The gene expression data of 17 tissues was obtained from the GTEx project for human and ENCODE project for mouse. The homologous genes between mouse and human were downloaded from the Mouse Genome Informatics (MGI) database [70]. For each homologous gene, the Z scores of TPM across the 17 tissues were calculated. If the difference in Z scores between human and mouse exceeded 1 in a given tissue, the tissue expression pattern of the homologous gene was considered altered.

The ChIP-seq profiles of HNF4A, HNF4G, FOXA1, and FOXA2 in the human liver were collected from ENCODE project and displayed on WashU Epigenome Browser. FIMO software [42] was used to identify the TFBS motifs in the consensus sequence of MER67A and the MER67A promoter of IL27 based on motif weigh matrix file from JASPAR [71].

## Declarations

### Ethics approval and consent to participate

Not applicable.

### Consent for publication

Not applicable.

### Availability of data and materials

Details about data analyzed in this study were included in the Methods section. The modified pipeline, together with annotations of TE-initiated transcripts produced in this study are available in (https://github.com/yunzhang1998/TEIRI).

### Author contributions

E.Y. conceived and designed the study. Y.Z., J.M.Z. and S.Y.H. performed the experiments. Y.Z., J.Q.S., X.Y.H., Y.Q.J., M.H.D. and C.Y.T. analyzed the data. Y.Z., J.Q.S., and E.Y. interpreted the data and wrote the manuscript. All authors approved the final version of the manuscript.

## Abbreviations

5’ RACE: 5’ rapid amplification of cDNA ends
ADME: Absorption, distribution, metabolism, and excretion
BH: Benjamini-Hochberg
CAGE: Cap analysis gene expression
CDS: Coding sequence
ChIP: chromatin immunoprecipitation
ENTEx: Epigenomes from four individuals
FANTOM: Functional annotation of the mammalian genome
FDR: False discovery rate
GEO: Gene expression omnibus
GRCh38: Genome reference consortium human build 38
GTEx: Genotype-Tissue Expression
HERV: Human endogenous retrovirus
hSSCs: Human adult spermatogonial stem cells
LINE: Long interspersed nuclear element
lncRNA: Long noncoding RNA
LTR: Long terminal repeat
MGI: Mouse Genome Informatics
ONT: Oxford Nanopore Technologies
PacBio: Pacific Biosciences
RAMPAGE: RNA annotation and mapping of promoters for the analysis of gene expression
SINE: Short interspersed nuclear element
TE: Transposable elements
TFBS: Transcription factor binding site
TSS: Transcription start site
UMAP: Uniform Manifold Approximation and Projection

## Acknowledgments

The Genotype-Tissue Expression (GTEx) Project was supported by the Common Fund of the Office of the Director of the National Institutes of Health, and by NCI, NHGRI, NHLBI, NIDA, NIMH, and NINDS. We acknowledge the FANTOM5 project, ENCODE Consortium and the ENCODE production laboratories.

## Additional files

**Additional file 1: Fig. S1-6.** Supplementary figure legends and supplementary figures (Fig. S1-S6, PDF).

**Additional file 2: Table S1-11.** Supplementary tables (Table S1-S11, XLSX).

## References

1. Britten RJ, Kohne DE. Repeated sequences in DNA. Hundreds of thousands of copies of DNA sequences have been incorporated into the genomes of higher organisms. Science. 1968 Aug 9;161(3841):529–40.

2. Mcclintock B. Controlling elements and the gene. Cold Spring Harb Symp Quant Biol. 1956;21:197–216.

3. Kapitonov VV, Jurka J. A universal classification of eukaryotic transposable elements implemented in Repbase. Nat Rev Genet. 2008 May;9(5):411–2; author reply 414.

4. Wells JN, Feschotte C. A Field Guide to Eukaryotic Transposable Elements. Annu Rev Genet. 2020 Sep 21;54:539.

5. Andrews G, Fan K, Pratt HE, Phalke N, Zoonomia Consortium§, Karlsson EK, et al. Mammalian evolution of human cis-regulatory elements and transcription factor binding sites. Science. 2023 Apr 28;380(6643):eabn7930.

6. Miao B, Fu S, Lyu C, Gontarz P, Wang T, Zhang B. Tissue-specific usage of transposable element-derived promoters in mouse development. Genome Biol. 2020 Sep 28;21(1):255.

7. Sun MA, Wolf G, Wang Y, Senft AD, Ralls S, Jin J, et al. Endogenous Retroviruses Drive Lineage-Specific Regulatory Evolution across Primate and Rodent Placentae. Mol Biol Evol. 2021 Oct 27;38(11):4992–5004.

8. Wu F, Liufu Z, Liu Y, Guo L, Wu J, Cao S, et al. Species-specific rewiring of definitive endoderm developmental gene activation via endogenous retroviruses through TET1-mediated demethylation. Cell Rep. 2022 Dec 13;41(11):111791.

9. Zhang Y, Li T, Preissl S, Amaral ML, Grinstein JD, Farah EN, et al. Transcriptionally active HERV-H retrotransposons demarcate topologically associating domains in human pluripotent stem cells. Nat Genet. 2019 Sep;51(9):1380–8.

10. Schrader L, Schmitz J. The impact of transposable elements in adaptive evolution. Mol Ecol. 2019 Mar;28(6):1537–49.

11. Deniz Ö, Frost JM, Branco MR. Regulation of transposable elements by DNA modifications. Nat Rev Genet. 2019 Jul;20(7):417–31.

12. Hollister JD, Gaut BS. Epigenetic silencing of transposable elements: a trade-off between reduced transposition and deleterious effects on neighboring gene expression. Genome Res. 2009 Aug;19(8):1419–28.

13. Mi S, Lee X, Li X, Veldman GM, Finnerty H, Racie L, et al. Syncytin is a captive retroviral envelope protein involved in human placental morphogenesis. Nature. 2000 Feb 17;403(6771):785–9.

14. Modzelewski AJ, Shao W, Chen J, Lee A, Qi X, Noon M, et al. A mouse-specific retrotransposon drives a conserved Cdk2ap1 isoform essential for development. Cell. 2021 Oct 28;184(22):5541–5558.e22.

15. Pasquesi GIM, Perry BW, Vandewege MW, Ruggiero RP, Schield DR, Castoe TA. Vertebrate Lineages Exhibit Diverse Patterns of Transposable Element Regulation and Expression across Tissues. Genome Biol Evol. 2020 May 1;12(5):506–21.

16. Tokuyama M, Kong Y, Song E, Jayewickreme T, Kang I, Iwasaki A. ERVmap analysis reveals genome-wide transcription of human endogenous retroviruses. Proc Natl Acad Sci U S A. 2018 Nov 19;115(50):12565.

17. Zhang XO, Gingeras TR, Weng Z. Genome-wide analysis of polymerase III–transcribed Alu elements suggests cell-type–specific enhancer function. Genome Res. 2019 Sep;29(9):1402.

18. Zhang F, Raabe CA, Cardoso-Moreira M, Brosius J, Kaessmann H, Schmitz J. ExoPLOT: Representation of alternative splicing in human tissues and developmental stages with transposed element (TE) involvement. Genomics. 2022 Jul;114(4):110434.

19. She J, Du M, Xu Z, Jin Y, Li Y, Zhang D, et al. The landscape of hervRNAs transcribed from human endogenous retroviruses across human body sites. Genome Biol. 2022 Nov 3;23(1):231.

20. Gu X, Wang M, Zhang XO. TE-TSS: an integrated data resource of human and mouse transposable element (TE)-derived transcription start site (TSS). Nucleic Acids Res. 2023 Nov 13;52(D1):D322.

21. Jang HS, Shah NM, Du AY, Dailey ZZ, Pehrsson EC, Godoy PM, et al. Transposable elements drive widespread expression of oncogenes in human cancers. Nat Genet. 2019 Apr;51(4):611–7.

22. Shah NM, Jang HJ, Liang Y, Maeng JH, Tzeng SC, Wu A, et al. Pan-cancer analysis identifies tumor-specific antigens derived from transposable elements. Nat Genet. 2023 Apr;55(4):631–9.

23. Carter TA, Singh M, Dumbović G, Chobirko JD, Rinn JL, Feschotte C. Mosaic cis-regulatory evolution drives transcriptional partitioning of HERVH endogenous retrovirus in the human embryo. eLife. 2022 Feb 18;11:e76257.

24. Mangoni D, Simi A, Lau P, Armaos A, Ansaloni F, Codino A, et al. LINE-1 regulates cortical development by acting as long non-coding RNAs. Nat Commun. 2023 Aug 17;14(1):4974.

25. Garza R, Atacho DAM, Adami A, Gerdes P, Vinod M, Hsieh P, et al. LINE-1 retrotransposons drive human neuronal transcriptome complexity and functional diversification. Sci Adv. 2023 Nov 3;9(44):eadh9543.

26. ENCODE Project Consortium. An integrated encyclopedia of DNA elements in the human genome. Nature. 2012 Sep 6;489(7414):57–74.

27. GTEx Consortium, Laboratory, Data Analysis &Coordinating Center (LDACC)—Analysis Working Group, Statistical Methods groups—Analysis Working Group, Enhancing GTEx (eGTEx) groups, NIH Common Fund, NIH/NCI, et al. Genetic effects on gene expression across human tissues. Nature. 2017 Oct 11;550(7675):204–13.

28. Lizio M, Harshbarger J, Shimoji H, Severin J, Kasukawa T, Sahin S, et al. Gateways to the FANTOM5 promoter level mammalian expression atlas. Genome Biol. 2015 Jan 5;16(1):22.

29. Wang F, Xu Y, Wang R, Zhang B, Smith N, Notaro A, et al. TEQUILA-seq: a versatile and low-cost method for targeted long-read RNA sequencing. Nat Commun. 2023 Aug 8;14(1):4760.

30. Storer J, Hubley R, Rosen J, Wheeler TJ, Smit AF. The Dfam community resource of transposable element families, sequence models, and genome annotations. Mob DNA. 2021 Jan 12;12(1):2.

31. Chuong EB, Elde NC, Feschotte C. Regulatory activities of transposable elements: from conflicts to benefits. Nat Rev Genet. 2016 Nov 21;18(2):71.

32. Xie M, Hong C, Zhang B, Lowdon RF, Xing X, Li D, et al. DNA hypomethylation within specific transposable element families associates with tissue-specific enhancer landscape. Nat Genet. 2013 Jul;45(7):836–41.

33. Marill J, Cresteil T, Lanotte M, Chabot GG. Identification of human cytochrome P450s involved in the formation of all-trans-retinoic acid principal metabolites. Mol Pharmacol. 2000 Dec;58(6):1341–8.

34. Ben-Porath I, Thomson MW, Carey VJ, Ge R, Bell GW, Regev A, et al. An embryonic stem cell-like gene expression signature in poorly differentiated aggressive human tumors. Nat Genet. 2008 May;40(5):499–507.

35. Guo J, Grow EJ, Yi C, Mlcochova H, Maher GJ, Lindskog C, et al. Chromatin and Single-Cell RNA-Seq Profiling Reveal Dynamic Signaling and Metabolic Transitions during Human Spermatogonial Stem Cell Development. Cell Stem Cell. 2017 Oct 5;21(4):533–546.e6.

36. Ito J, Sugimoto R, Nakaoka H, Yamada S, Kimura T, Hayano T, et al. Systematic identification and characterization of regulatory elements derived from human endogenous retroviruses. PLoS Genet. 2017 Jul;13(7):e1006883.

37. Hayashi Y, Caboni L, Das D, Yumoto F, Clayton T, Deller MC, et al. Structure-based discovery of NANOG variant with enhanced properties to promote self-renewal and reprogramming of pluripotent stem cells. Proc Natl Acad Sci U S A. 2015 Apr 14;112(15):4666–71.

38. Li L, Wang Y, Torkelson JL, Shankar G, Pattison JM, Zhen HH, et al. TFAP2C– and p63-Dependent Networks Sequentially Rearrange Chromatin Landscapes to Drive Human Epidermal Lineage Commitment. Cell Stem Cell. 2019 Feb 7;24(2):271–284.e8.

39. Zoonomia Consortium. A comparative genomics multitool for scientific discovery and conservation. Nature. 2020 Nov;587(7833):240–5.

40. Kryuchkova-Mostacci N, Robinson-Rechavi M. A benchmark of gene expression tissue-specificity metrics. Brief Bioinform. 2017 Mar 1;18(2):205–14.

41. Wang Q, Li D, Cao G, Shi Q, Zhu J, Zhang M, et al. IL-27 signalling promotes adipocyte thermogenesis and energy expenditure. Nature. 2021 Dec;600(7888):314–8.

42. Bailey TL, Boden M, Buske FA, Frith M, Grant CE, Clementi L, et al. MEME SUITE: tools for motif discovery and searching. Nucleic Acids Res. 2009 Jul;37(Web Server issue):W202–208.

43. Fueyo R, Judd J, Feschotte C, Wysocka J. Roles of transposable elements in the regulation of mammalian transcription. Nat Rev Mol Cell Biol. 2022 Jul;23(7):481–97.

44. Oliveira DS, Fablet M, Larue A, Vallier A, Carareto CMA, Rebollo R, et al. ChimeraTE: a pipeline to detect chimeric transcripts derived from genes and transposable elements. Nucleic Acids Res. 2023 Oct 13;51(18):9764–84.

45. Senft AD, Macfarlan TS. Transposable elements shape the evolution of mammalian development. Nat Rev Genet. 2021 Nov;22(11):691–711.

46. Stark R, Grzelak M, Hadfield J. RNA sequencing: the teenage years. Nat Rev Genet. 2019 Nov;20(11):631–56.

47. Batut P, Dobin A, Plessy C, Carninci P, Gingeras TR. High-fidelity promoter profiling reveals widespread alternative promoter usage and transposon-driven developmental gene expression. Genome Res. 2013 Jan;23(1):169–80.

48. Garalde DR, Snell EA, Jachimowicz D, Sipos B, Lloyd JH, Bruce M, et al. Highly parallel direct RNA sequencing on an array of nanopores. Nat Methods. 2018 Mar;15(3):201–6.

49. Wenger AM, Peluso P, Rowell WJ, Chang PC, Hall RJ, Concepcion GT, et al. Accurate circular consensus long-read sequencing improves variant detection and assembly of a human genome. Nat Biotechnol. 2019 Oct;37(10):1155–62.

50. Pardo-Palacios FJ, Wang D, Reese F, Diekhans M, Carbonell-Sala S, Williams B, et al. Systematic assessment of long-read RNA-seq methods for transcript identification and quantification. Nat Methods. 2024 Jul;21(7):1349–63.

51. Carbonell-Sala S, Perteghella T, Lagarde J, Nishiyori H, Palumbo E, Arnan C, et al. CapTrap-seq: a platform-agnostic and quantitative approach for high-fidelity full-length RNA sequencing. Nat Commun. 2024 Jun 27;15(1):5278.

52. Bourque G, Leong B, Vega VB, Chen X, Lee YL, Srinivasan KG, et al. Evolution of the mammalian transcription factor binding repertoire via transposable elements. Genome Res. 2008 Nov;18(11):1752–62.

53. Chuong EB, Elde NC, Feschotte C. Regulatory evolution of innate immunity through co-option of endogenous retroviruses. Science. 2016 Mar 4;351(6277):1083–7.

54. Uhlén M, Fagerberg L, Hallström BM, Lindskog C, Oksvold P, Mardinoglu A, et al. Proteomics. Tissue-based map of the human proteome. Science. 2015 Jan 23;347(6220):1260419.

55. Dobin A, Davis CA, Schlesinger F, Drenkow J, Zaleski C, Jha S, et al. STAR: ultrafast universal RNA-seq aligner. Bioinformatics. 2012 Oct 25;29(1):15.

56. Li H. New strategies to improve minimap2 alignment accuracy. Bioinforma Oxf Engl. 2021 Dec 7;37(23):4572–4.

57. Kovaka S, Zimin AV, Pertea GM, Razaghi R, Salzberg SL, Pertea M. Transcriptome assembly from long-read RNA-seq alignments with StringTie2. Genome Biol. 2019 Dec 16;20(1):278.

58. Harrow J, Frankish A, Gonzalez JM, Tapanari E, Diekhans M, Kokocinski F, et al. GENCODE: the reference human genome annotation for The ENCODE Project. Genome Res. 2012 Sep;22(9):1760–74.

59. Tang AD, Soulette CM, van Baren MJ, Hart K, Hrabeta-Robinson E, Wu CJ, et al. Full-length transcript characterization of SF3B1 mutation in chronic lymphocytic leukemia reveals downregulation of retained introns. Nat Commun. 2020 Mar 18;11(1):1438.

60. Quinlan AR, Hall IM. BEDTools: a flexible suite of utilities for comparing genomic features. Bioinforma Oxf Engl. 2010 Mar 15;26(6):841–2.

61. Ramírez F, Ryan DP, Grüning B, Bhardwaj V, Kilpert F, Richter AS, et al. deepTools2: a next generation web server for deep-sequencing data analysis. Nucleic Acids Res. 2016 Jul 8;44(W1):W160–165.

62. Zhou X, Wang T. Using the Wash U Epigenome Browser to examine genome-wide sequencing data. Curr Protoc Bioinforma. 2012 Dec; Chapter 10:10.10.1-10.10.14.

63. Pertea G, Pertea M. GFF Utilities: GffRead and GffCompare. F1000Research. 2020;9:ISCB Comm J-304.

64. Kang YJ, Yang DC, Kong L, Hou M, Meng YQ, Wei L, et al. CPC2: a fast and accurate coding potential calculator based on sequence intrinsic features. Nucleic Acids Res. 2017 Jul 3;45(W1):W12–6.

65. Camacho C, Coulouris G, Avagyan V, Ma N, Papadopoulos J, Bealer K, et al. BLAST+: architecture and applications. BMC Bioinformatics. 2009 Dec 15;10:421.

66. McInnes L, Healy J, Melville J. UMAP: Uniform Manifold Approximation and Projection for Dimension Reduction [Internet]. arXiv; 2020 [cited 2024 Oct 23]. Available from: http://arxiv.org/abs/1802.03426

67. Yanai I, Benjamin H, Shmoish M, Chalifa-Caspi V, Shklar M, Ophir R, et al. Genome-wide midrange transcription profiles reveal expression level relationships in human tissue specification. Bioinforma Oxf Engl. 2005 Mar 1;21(5):650–9.

68. Yu G, He QY. ReactomePA: an R/Bioconductor package for reactome pathway analysis and visualization. Mol Biosyst. 2016 Feb;12(2):477–9.

69. Katoh K, Standley DM. MAFFT multiple sequence alignment software version 7: improvements in performance and usability. Mol Biol Evol. 2013 Apr;30(4):772–80.

70. Baldarelli RM, Smith CL, Ringwald M, Richardson JE, Bult CJ, Mouse Genome Informatics Group. Mouse Genome Informatics: an integrated knowledgebase system for the laboratory mouse. Genetics. 2024 May 7;227(1):iyae031.

71. Rauluseviciute I, Riudavets-Puig R, Blanc-Mathieu R, Castro-Mondragon JA, Ferenc K, Kumar V, et al. JASPAR 2024: 20th anniversary of the open-access database of transcription factor binding profiles. Nucleic Acids Res. 2024 Jan 5;52(D1):D174–82.

